# From high protection to lethal effect: diverse outcomes of immunization against invasive candidiasis with different *Candida albicans* extracellular vesicles

**DOI:** 10.1101/2024.11.21.624473

**Authors:** Raquel Martínez-López, Gloria Molero, Claudia Parra-Giraldo, Matías Cabeza, Guillermo Castejón, Carmen García-Duran, Felipe Clemente, María Luisa Hernáez, Concha Gil, Lucía Monteoliva

## Abstract

Extracellular vesicles (EVs) from *Candida albicans* can elicit immune responses, positioning them as promising acellular vaccine candidates. We characterized EVs from an avirulent *C. albicans* cell wall mutant (*ecm33Δ*) and evaluated their protective potential against invasive candidiasis. EVs from the yeast (YEVs) and hyphal (HEVs) forms of the SC5314 wild-type strain were also tested, yielding high survival rates with SC5314 YEVs (91%) and *ecm33* YEVs immunization (64%). Surprisingly, HEV immunization showed a dual effect, resulting in 36% protection but also causing premature death in some mice. Proteomic analyses revealed distinct profiles among the top 100 proteins in the different EVs which may explain these effects: a shared core of 50 immunogenic proteins such as Pgk1, Cdc19, and Fba1; unique, relevant immunogenic proteins in SC5314 YEVs, and proteins linked to pathogenesis, like Ece1 in SC5314 HEVs. Sera from SC5314 YEVs-immunized mice showed the highest IgG2a titers and moderate IL-17, IFN-γ, and TNF-α levels, indicating the importance of both humoral and cellular responses for protection. These findings highlight the distinct immunogenic properties of *C. albicans* EVs, suggesting their potential in acellular vaccine development while emphasizing the need to carefully evaluate pathogenic risks associated with certain EVs.

## 1. Introduction

Invasive fungal infections mainly affect immunocompromised patients and their incidence is underestimated [1]. In 2022, the World Health Organization (WHO) released the first-ever list of health-threatening fungi, which includes six *Candida* species: *Candida albicans, Candida auris, Candida parapsilosis, Candida tropicalis, Nakaseomyces glabrata* (*Candida glabrata*) *and Pichia kudriavzeveii* (*Candida krusei*) [2]. Invasive candidiasis (IC) represents a significant threat to immunocompromised individuals. More than 1,5 million people worldwide are estimated to be affected by IC annually, with a mortality rate of 64% [1]. *C. albicans* is the most frequently isolated cause [3]. *C. auris* also worries the medical community due to its persistence in the ICUs environment and its resistance to the usual antifungal treatments. The difficulties of managing patients with IC are related to late diagnosis, the limitations of the antifungal treatment with only a few drugs available, and the increase in antifungal resistance [4].

*C. albicans* is a commensal polymorphic fungus that causes different types of infections in immunocompromised hosts, with IC being the most serious one. *C. albicans* can grow in a yeast form or as hyphae, among other cell morphologies, and it is able to form biofilms. Yeast and hyphae have different roles during the infection, and biofilms are a relevant medical problem because they grow on catheters or prostheses and are more resistant to antifungal drugs and host defenses [3-5]. Therefore, studies to design new strategies to prevent and treat IC are of interest.

Vaccination stands out as a promising approach to fight IC. Many attempts have been made by different groups to design a vaccine against candidiasis [6-8]. However, developing a safe and effective vaccine remains a challenge [6, 8]. The groups that have tested vaccines with live cells of mutants with reduced virulence, such as *hog1* [9, 10], *ecm33* [11], or *nrg1* [12], have reported high levels of protection in mice. However, a live vaccine is not advisable for many target groups, such as pregnant women and immunocompromised individuals; thus, our studies have been focused on identifying antigenic proteins with a possible role in protection. Based on previous studies of diagnostic and prognostic biomarkers of IC identified in sera from human patients [13-19], we investigated the highly immunogenic protein Bgl2 for vaccination, achieving 25 % of protection [20]. Other trials with single proteins have been published, including Hyr1 [21], Trx1[22], and several moonlighting *Candida* proteins [23]. However, the only vaccines reaching Phase 2 clinical trials have been the ones based on a single protein, either Als3 or Sap2, against vulvovaginal recurrent candidiasis (VVRC). The Als3 vaccine has proven the protection against *C. albicans* [24], *C. auris* [25] and *Staphylococcus aureus* (systemic infection) [26, 27]. A recent study used Cht3 as the antigen and lipid particles as the delivery system [28]. Extracellular vesicles (EVs) are membrane-bound particles released by cells, including bacterial and human cells, into the extracellular space. They can carry various cargo, including proteins, lipids, and nucleic acids and enzymes. EVs include nanosized exosomes (70−150 nm), derived from the endosomal system, or microvesicles (100−1000 nm), produced by outward budding of the plasma membrane. EVs are now considered an important mechanism of intercellular communication, allowing cells to exchange proteins, lipids, and genetic material. There has been a sharp increase in scientific interest in the physiological and pathological functions of fungal and human EVs, as they have potential biomedical applications [29, 30].

The use of EVs as a vaccine platform offers several advantages, including mimicking natural infection, potential targeted delivery of antigens to antigen-presenting cells, and the induction of both innate and adaptive immune responses. In this regard, there are precedents for EVs in vaccines, such as the case of the anti-meningococcal vaccine Bexsero, used against *Neisseria meningitidis* serogroup B invasive disease [31], which is composed of outer membrane vesicles and some immunogenic proteins.

In fungi, EVs are involved in pathogenesis and immune evasion, and may be associated with cell-cell communication [32, 33]. As fungal EVs may play a pivotal role in the establishment of fungal infections and can alter the infection process, they are possible targets for new antifungal agents and potential candidates for vaccine development [34]. These EVs may contain various fungal antigens, including cell wall components and virulence factors, as well as host-derived molecules enhancing the immune response [20, 35]. EVs have also been used as vehicles for proteins that can have antigenic potential. Vargas et al. (2020) used *C. albicans* EVs as vaccine and achieved great protection against a *C. albicans* peritoneal invasive infection [36].

Our group assessed the protective efficacy of alive cells of the *C. albicans ecm33Δ* mutant. This mutant is characterized by an altered cell wall with an increased outermost protein layer and has demonstrated promising results in BALB/c mice, providing effective protection in a systemic candidiasis mouse model [11]. In another study, we investigated the proteins secreted by this mutant growing in synthetic minimal medium, categorizing them into a vesicle-free secretome and EVs [37]. Notably, 98 proteins were common to both fractions, predominantly representing metabolic and cell wall-related proteins, highlighting a modified pattern of protein secretion through the classical pathway and distinct characteristics in EVs formation with respect to the wild-type SC5314 strain [37].

Moreover, due to the importance of the yeast-to-hypha transition in virulence and the establishment of IC, we analyzed the protein composition and cargo of EVs derived from yeast and hyphal forms of *C. albicans* [35]. The characterization of EVs secreted by *C. albicans* hyphal cells (HEVs) revealed distinctive features compared to EVs from the yeast form (YEVs). YEVs are rich in cell wall proteins and have demonstrated the ability to rescue the sensitivity of a cell wall mutant against calcofluor white. Conversely, HEVs exhibited greater protein diversity and a higher proportion of virulence-associated proteins. Immunogenic assessments revealed that both YEVs and HEVs elicited immune reactivity with human sera from IC patients, however, HEVs exhibited cytotoxic effects on human macrophages and induced the release of tumor necrosis factor alpha (TNF-α) [35]. This comprehensive analysis highlighted the distinct roles of YEVs and HEVs, with the former playing an important role in cell wall maintenance and the latter potentially associated with virulence. These findings showcase their potential as valuable targets for vaccine development and insights into novel diagnostic markers and treatment strategies against *C. albicans* infections.

Our ongoing research focuses on studying EVs derived from different strains and morphological forms of *C. albicans* and assessing their potential for vaccination in a murine model of IC.

## 2. Results

### 2.1. Characterization of EVs secreted by ecm33Δ mutant growing in a rich medium

YEVs secreted by the *ecm33*Δ cell wall mutant (*ecm33* YEVs) were isolated and morphologically characterized. These vesicles were compared with YEVs derived from the SC5314 parental wild-type strain (SC5314 YEVs). Both were cultured under identical conditions (16 h of growth in YPD at 37°C and 180 rpm). Electron microscope images revealed typical spherical structures with a denser bilayer. Nanoparticle tracking analysis (NTA) revealed consistent vesicle sizes of 100 - 200 nm for both strains (Figure 1).

**Figure 1.**
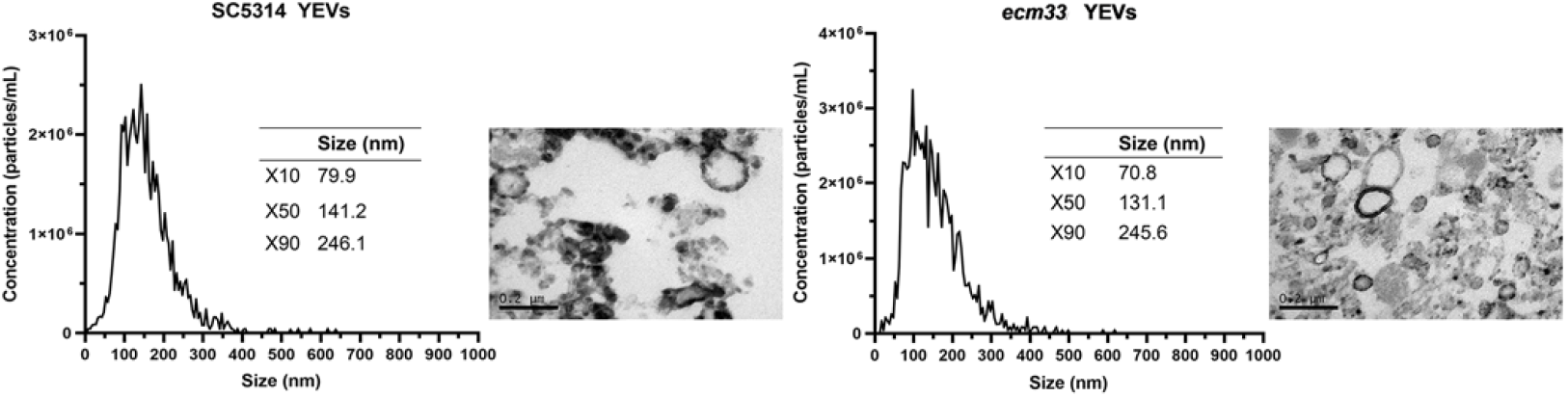
NTA analysis of SC5314 YEVs and *ecm33* YEVs. TEM images showing spherical electron-dense bilayered structures, typical of extracellular vesicles, are provided for visual reference. The X10, X50 and X90 indicate a 10%, 50% or 90% of EVs respectively with the specified size or smaller.

To compare the differences in the protein cargo of the EVs secreted by the wild-type and the mutant strains, we carried out a proteomic study. Three biological replicates of each type of EVs were analyzed. We identified a total of 1443 proteins: 1219 (Table S1) and 1333 (Table S2) corresponding to SC5314 YEVs and *ecm33* YEVs, respectively, and 1109 proteins in common (Figure 2a). The Gene Ontology (GO) enrichment analysis of the different protein sets revealed a higher enrichment of cell wall and cell surface proteins in the SC5314 YEVs compared to *ecm33* YEVs (Figure 2a).

**Figure 2.**
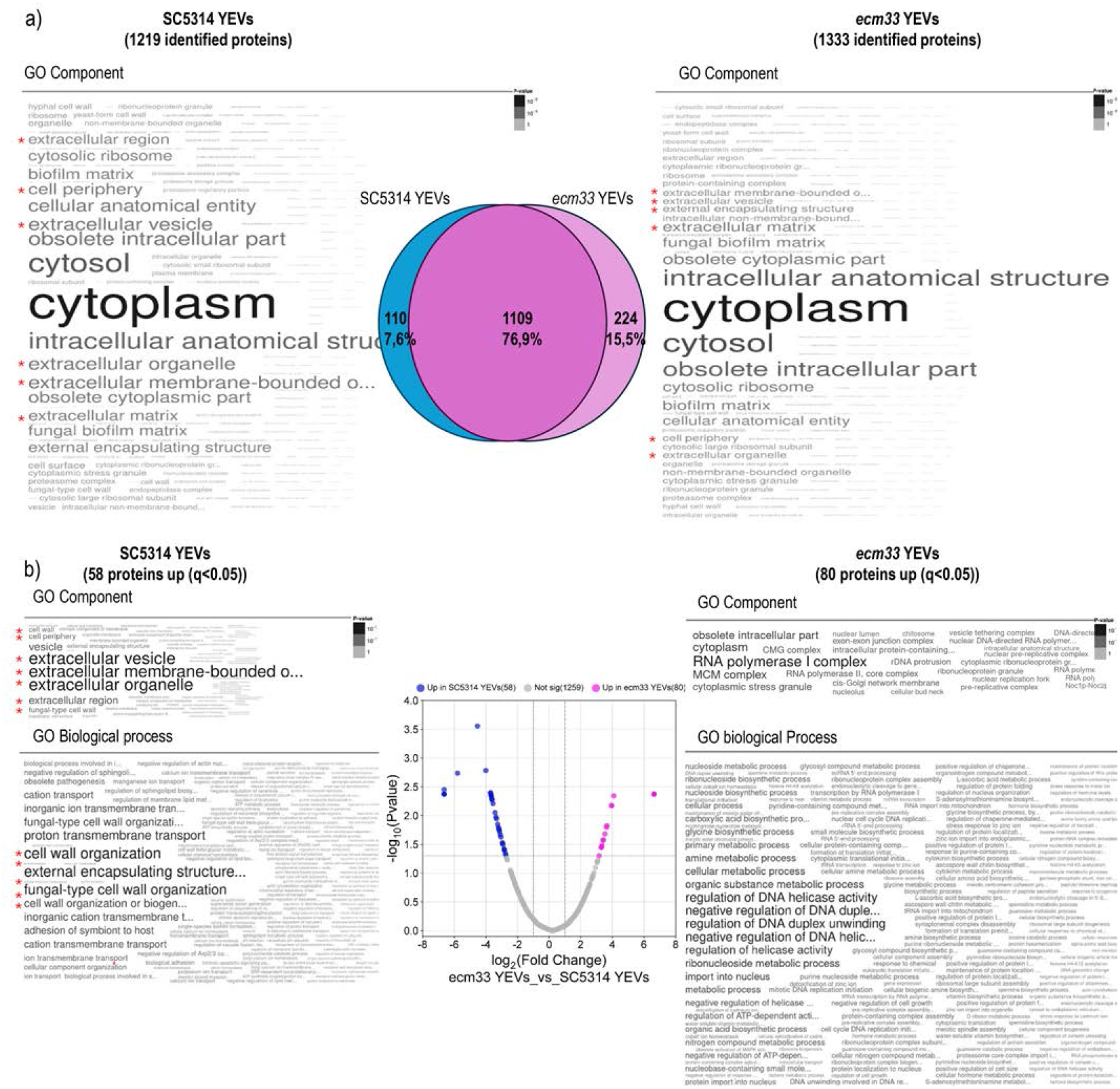
**a)** Venn diagram illustrating the proportion of proteins identified in both type of EVs or exclusively in SC5314 YEVS or *ecm33* YEVs, along with an enrichment analysis based on Gene Ontology (GO) cellular component from the Candida Genome Database. b) Volcano plot (SRplot software) depicting quantified proteins with a q value <0.05, alongside an enrichment analysis based on GO considering both cellular component and biological process. The red asterisks indicate cellular components or biological processes related to the cell wall or cell surface.

In the label-free proteomic analysis, a total of 1397 proteins were quantified (Table S3), 58 of which were more abundant in SC5314 YEVs and 80 in *ecm33* YEVs (q value <0.05; Fig. 2b, Table 1 and Table S4). Considering the differentially abundant proteins in both types of EVs and the GO enrichment analysis of both cellular component and biological process, a clear enrichment of cell surface proteins was confirmed in SC5314 YEVs (Fig. 2b). In addition, a set of four proteins (Sap2, Sap10, Ecm33, and Als2), related to host pathogenesis [38-41] was found exclusively in SC5314 YEVs (Table 1 and Table S4).

**Table 1.** List of proteins with higher abundance (or exclusively detected) in SC5314 YEVs and *ecm33* YEVs, with a q-value < 0.05. Proteins are grouped according to GO terms from the Candida Genome Database.

| More abundant in SC5314 YEVs (58) |  | More abundant in *ecm33* YEVs (80) |  |
| --- | --- | --- | --- |
| Exclusively detected | Significantly more abundant | Exclusively detected | Significantly more abundant |
| **Extracellular region (16)** |  | **Nucleus (18)** |  |
| Als2, Ecm33, Gap4, Hgt6,<br>Lip4, Rbe1, Sap10, Sap2,<br>Sap7 | Bbc1, Nce102, Pga4, Sun41,<br>Tos1, Xog1, Sap8 | Blm3, Orf19.6340, Orf19.3778,<br>Orf19.6665, Orf19.3477, Cbk1,<br>Cdc5, Cdc54, Ecm29, Hat1,<br>Kap120, Mak21, Mcm2,<br>Rpa190, Rpb8, Rpp1, Sda1 | Prp24 |
| **Plasma membrane (10)** |  | **Membrane (9)** |  |
| Orf19.7670, Mrs7, Pmr1,<br>Cox1 | Orf19.409, Rta2, Orf19.4706, _<br>Atp6, Fmp45, Atp7 | Orf19.3307, Erg26, Erg28, Ole1,<br>Osh2, Zrc1, Vma11 | Chs2, Pst1 |
| **Endomembrane system (11)** |  | **Endomembrane system (6)** |  |
| Sla1, Orf19.6951,<br>Orf19.3128, Orf19.6552 | Ire1, Orf.2533.1, Ypt72, Orm1,<br>Orf19.3083, Orf19.6804, Cup5, | Arf1, Orf19.1994, Orf19.5817,<br>Cog4, Exo70 | Emp70 |
| **Mitochondrion (7)** |  | **Mitochondrion (5)** |  |
| Orf19.4932, Orf19.5296,<br>Orf19.7485, Mrps9,<br>Orf19.1956 | Rad51, Orf19.3154 | Bat21, Mrpl19, Ste23, Tom20 | Ptc2 |
| **Other (8)** |  | **Other (14)** |  |
| Orf19.1965, Gyp7, Tra1,<br>Orf19.5184 | Orf19.4758, Orf19.3792,<br>Orf19.6119, Orf19.1769 | Orf19.4529, orf19.864,<br>orf19.4030, Cgr1, Orf19.6305,<br>Gcd7, Pbs2, Sgt2, Tif11, Zta1 | Orf19.7109, Drg1,<br>Rpl42, Slk19 |
| **Cellular component unknown (6)** |  | **Cellular component unknown (28)** |  |
| Orf19.4061,<br>Orf19.1433, Orf19.163 | Orf19.1152, Orf19.5686, Pho8 | Bio32, Orf19.3309, Orf19.3679,<br>Orf19.2966, Orf19.2663,<br>Orf19.4846, Orf19.2269,<br>Orf19.1723, Orf19.5921,<br>Orf19.5038, Orf19.1249,<br>Orf19.4220, Orf19.1124,<br>Orf19.3219, Orf19.4043,<br>Orf19.5763, Orf19.6463, Cek2,<br>Hal22, Rki1, Spe2, Tif3,<br>Orf19.1864, Orf19.1078, Ura5 | orf19.597, Pnp1,<br>Pwp1 |

### 2.2. Immunization assays with EVs

We studied the optimal conditions to test the effect of the immunization with *ecm33* YEVs. High doses of 10 or 20 µg were used as the first approach (Figure S1). Mice were immunized by subcutaneous injection in two doses with a 2-week interval (Figure S1a). To determine the effect of including an adjuvant in the immunization formulation, another group of mice was immunized with 20 µg of *ecm33* YEVs plus aluminum adjuvant (1:2). Notably, prior to the administration of the second dose of EVs, an inflammatory nodule was observed at the site of the first dose in this later group. For this reason, the aluminum adjuvant proportion was reduced to 1:3 for the second immunization. Two control groups inoculated with PBS or PBS with adjuvant were also included. For the subsequent infection, a lethal intravenous injection of 1*x*106 cells was administered via the tail vein, similar to our previous successful vaccination assay with alive *ecm33*Δ mutant cells [11] (Figure S1a).

This first experiment resulted in an increased mortality rate post-infection in mice immunized with both doses of *ecm33* YEVs without adjuvant (Figure S1b), even surpassing that of the PBS control group. However, the 20µg dose of *ecm33* YEVs formulated with aluminum adjuvant provided a certain level of protection, with a 20% of survival rate. (Figure S1b).

In the light of these results, a reduction in both the vesicle dosage, to 5 µg, and the aluminum adjuvant, to a 1:3 proportion, were selected for the second immunization assay. In this assay, considering the significant differences in protein composition observed between *ecm33* YEVs and SC5314 YEVs, we assessed the immunization potential of both types of EVs. In addition, mice immunized with alive cells of the *C. albicans ecm33*Δ mutant were included to compare the immunization capacity of EVs with our previous results [11] (Figure 3a). Results showed a significant delay in mortality with respect to the control adjuvant group for both types of EVs (Figure 3b). The control group immunized with alive *ecm33*Δ cells reproduced the high level of protection published previously (Figure 3b), validating the new results.

**Figure 3.**
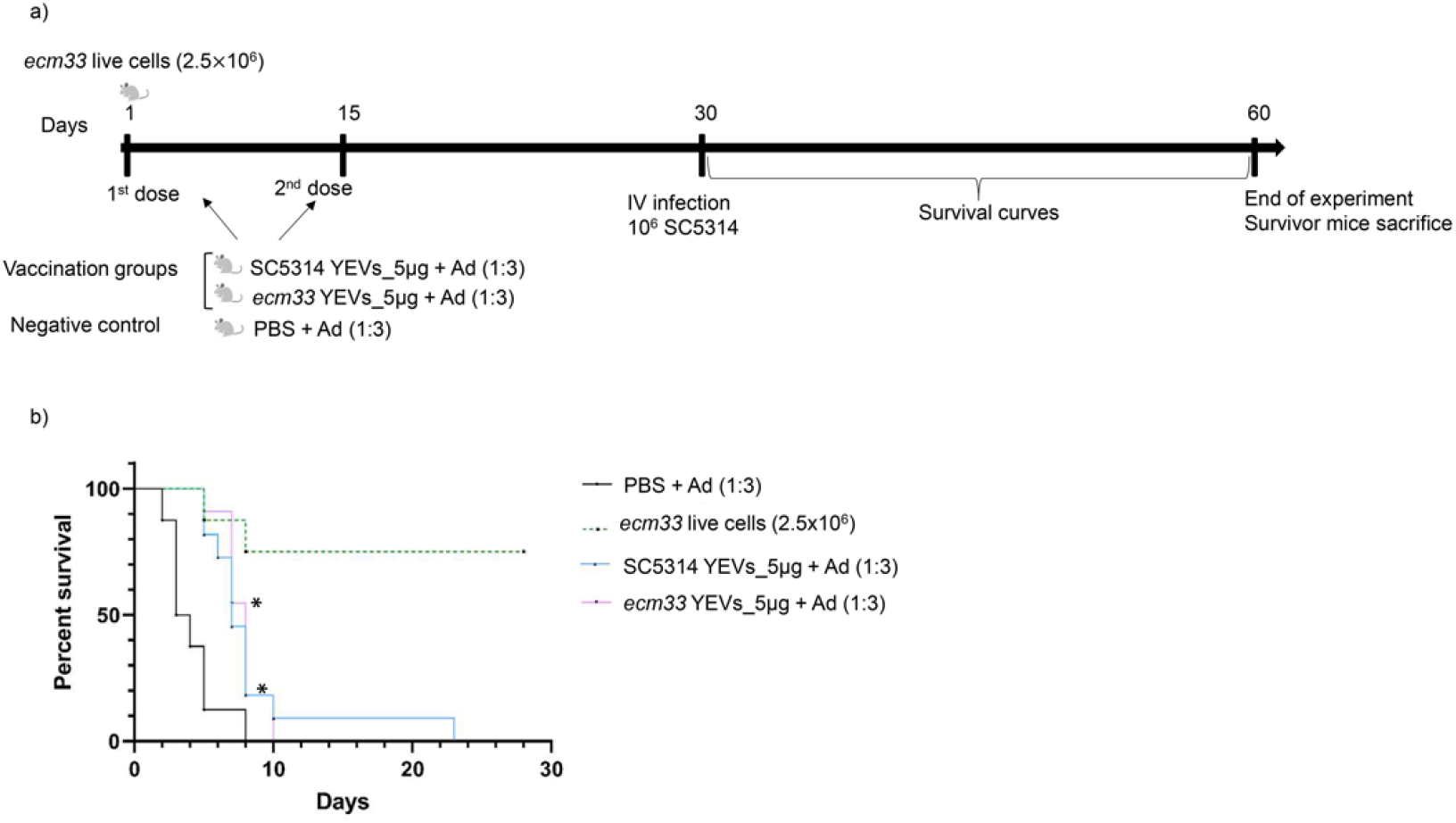
Vaccination schedule (a) and survival curves (b) of a murine model of IC showing the protective effect achieved with immunization with SC5314 YEVs and *ecm33* YEVs. Mice were immunized with different doses of EVs with adjuvant (Ad) and subsequently challenged with an intravenous lethal dose of SC5314 (1*x*10^6^ cells). Vaccination with live cells (2.5*x*10^6^ cells, one dose) of the completely avirulent *ecm33Δ* mutant was included as a positive vaccination control (a and b). * Mantel-cox test p <0.05

Immunization studies with acellular vaccines typically use a lower infective dose of *C. albicans*. Thus, a third experiment was performed with 5 *x* 10^5^cells/mL of parental strain SC5314 as the infection dosage (Figure 4a). As a second objective of this study was to assess the differences in the protective ability of the EVs secreted by the two morphological forms of *C. albicans* (yeast or hyphae), the EVs secreted by *C. albicans* hypha (SC5314 HEVs) were also included (Figure 4a). As in the previous experiments, mice were immunized by subcutaneous injection with two doses of 5 µg *ecm33* YEVs, SC5314 YEVs, or SC5314 HEVs (Figure 4a) plus the aluminum adjuvant (1:3). As shown in Figure 4b the survival rate of the mice immunized with SC5314 YEVs was 90%, whereas 100% of the mice from the control group died. On the other hand, immunization of mice with *ecm33* YEVs induced a lower level of protection (64%). Regarding the group of mice immunized with SC5314 HEVs, during the first week after infection, the survival rate was unexpectedly lower than that of the control group, though 36% of the mice remained alive at the end of the experiment (Figure 4b).

**Figure 4.**
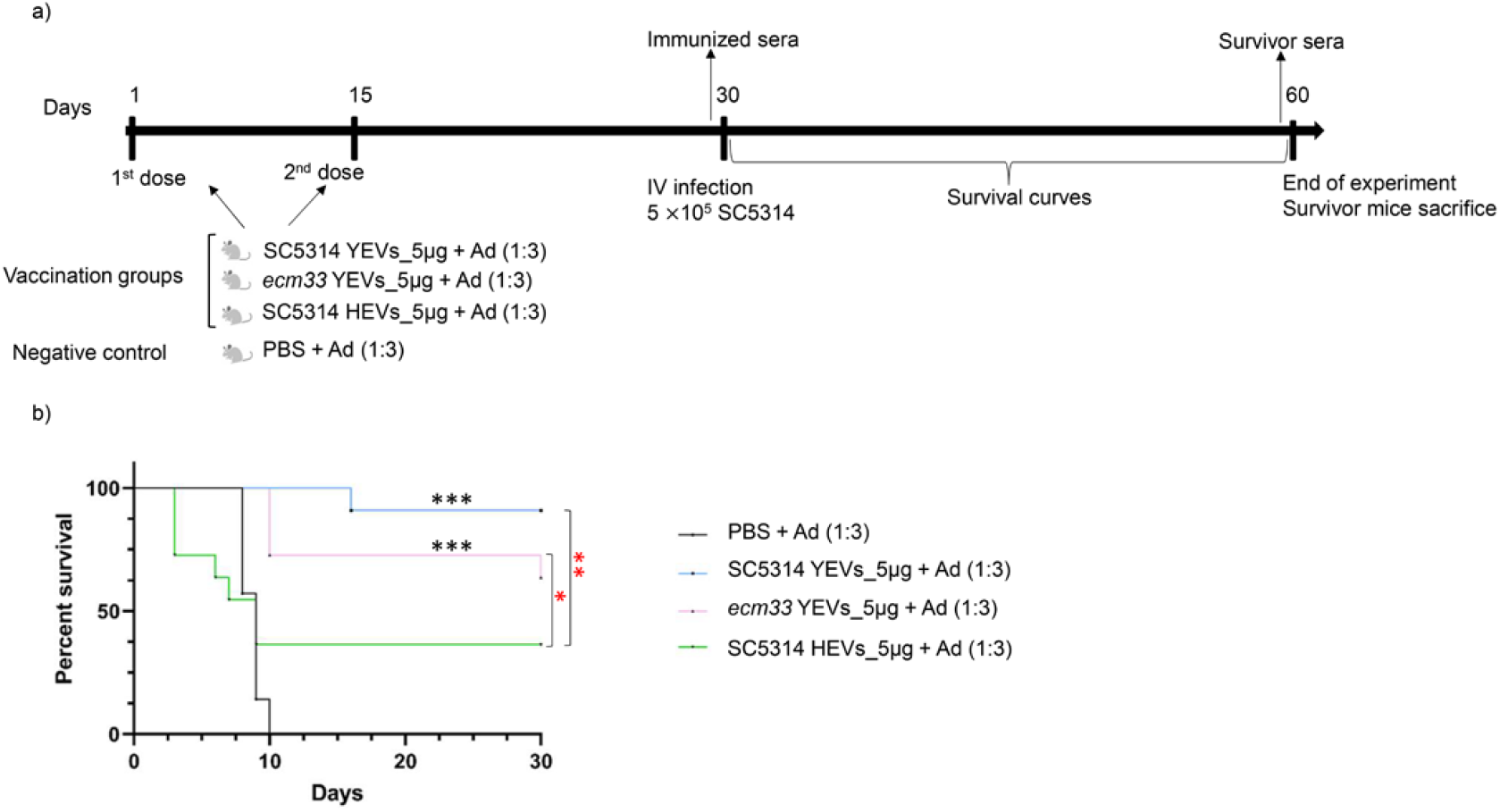
Vaccination schedule (a) and survival curves (b) of a mouse model of invasive candidiasis showing the protective effect of immunization with different *C. albicans* EVs. Mice were immunized with different doses of EVs with adjuvant (Ad) and subsequently challenged with an intravenous lethal dose of SC5314 (5×105 cells). Mantel-cox test * p <0.05, ** p <0.005, *** p <0.0005 (black asterisks represent statistically significance between the control group and the different EVs-immunized group. Red asterisks represent statistically significance in the protection acquired between the different EVs-immunized groups).

### 2.3. Analysis and comparison of the protein load of each of the three types of vesicles used for immunization

To understand why these EVs obtained from different *C. albicans* strains or morphological forms elicit such different responses against systemic candidiasis, we performed a comparative analysis of the protein composition and the serological response induced by each of the three types of EVs.

The label-free comparative analysis of the protein cargo of both the *ecm33* YEVs and SC5314 YEVs has already been described. In addition, a proteomic analysis of the SC5314 HEVs was published previously [35]. As the SC5314 HEVs analysis was not carried out simultaneously with the proteomic analysis currently presented, we could not perform a label-free analysis with the three types of EVs. Thus, we used the relative abundance of each protein within each sample based on the calculation of the Normalized Spectral Abundance Factor (NSAF). The NSAFs values, which are calculated considering the number of matched peptide spectra (PSMs) and the molecular weight, were used to conduct a clustering heatmap analysis with Rstudio (Figure 5a).

**Figure 5.**
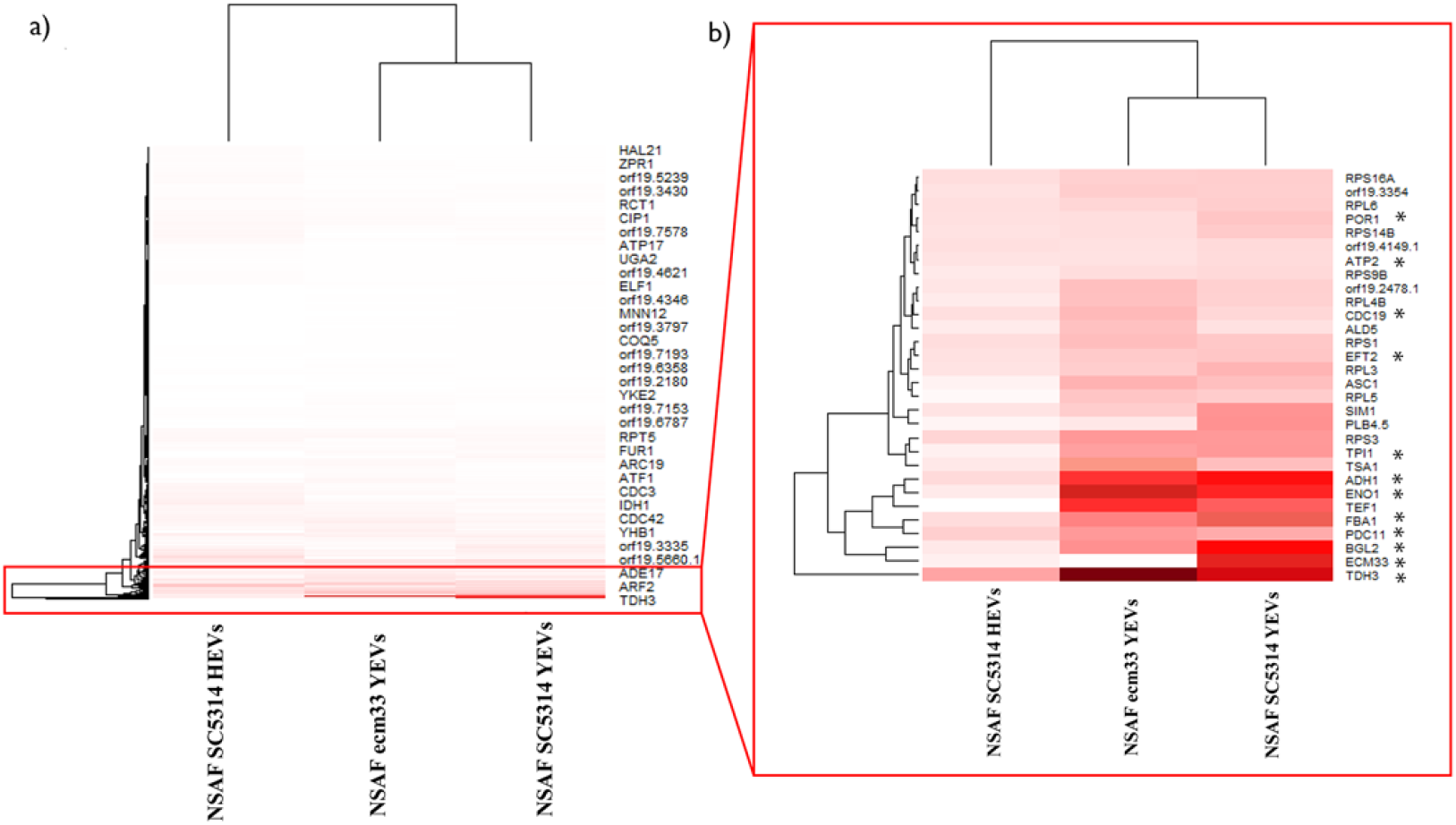
a) Hierarchical heat map depicting the relative abundance of each protein across the three different types of EVs, with darker shades of red indicating higher relative abundance (measured by NSAF) (Protein names shown represent 1 out of every 22 proteins for clarity) b) Zoomed-in view on the region of the heatmap with a higher abundance in cell surface proteins in SC5314 YEVs. Proteins describe as immunogenic in th Candida genome databased (CGD) are marked with an asterisk.

This analysis showed that the protein composition of the *ecm33* and SC5314 YEVs clustered together, whereas the protein profile of the SC5214 HEVs appeared separately in the dendrogram (Figure 5a). These differences in the protein composition correlate with the results of the vaccination experiment as the HEVs had the most different results (Figure 4 b). As previously observed, the proteins found in higher proportions in SC5314 YEVs corresponded to cell surface proteins, many of which have already been reported to be immunogenic (Figure 5b). These proteins were present in lower proportions in *ecm33* YEVs, and at even lower proportions in SC5314 HEVs (Figure 5b).

Focusing on the proteins most abundantly present in each vesicle type (top 100), all three vesicle types shared a core set of 50 proteins (Figure 6). Additionally, exclusive proteins were identified among the top 100 of each vesicle type: 35 unique proteins in SC5314 HEVs, 16 in SC5314 YEVs, and 19 in *ecm33* YEVs. The shared core includes several proteins previously identified as immunogenic, which may help explain why all three types demonstrated a certain degree of protective capacity. Notably, only the HEVs contained 7 proteins implicated in fungal virulence within their set of most abundant proteins (Arf1, Ece1, Hsp104, Phr1, Sap5, Sap6 and Ypt1), with Ece1—the precursor protein of candidalysin—standing out due to its direct association with Candida virulence. Among the proteins identified as most abundant exclusively in SC5314 YEVs, there are notable examples known for their strong immunogenic potential, such as Mp65, Mdh1 and Ecm33. The *ecm33* YEVs also include highly abundant proteins with immunogenic potential, such as Tkl1, Ilv5, and Ahp1.

**Figure 6.**
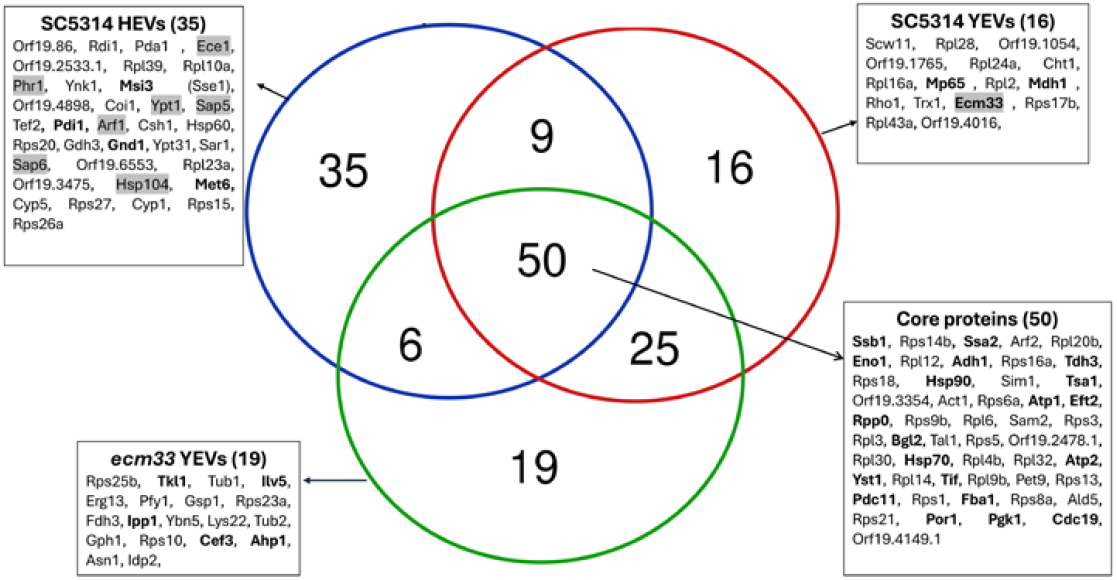
Venn diagram illustrating the top 100 most abundant proteins in each type of extracellular vesicle (EV). Proteins previously reported as immunogenic in other studies are highlighted in bold. Proteins associated with virulence are shaded.

### 2.4. Analysis of antibody titers in the sera of immunized mice

We conducted ELISAs with pooled sera obtained after EVs inmunization and prior to infection from mice in each of the groups used in the vaccination experiment shown in Figure 4 to study the IgG and the IgG2a antibody titers. As the proteins in the EVs used for immunization include cell surface proteins in an important proportion and it is difficult to obtain them in cytoplasmic cell extracts, three different protein samples were used for the determinations: proteins from SC5314 YEVs, proteins from HEVs, and a total yeast cytoplasmic extract of SC5314 strain obtained through mechanical disruption. As shown in Figure 7, all pooled sera had similar IgG titers against YEVs or HEVs proteins, while lower antibody titers were achieved against the cytoplasmic extracts for all sera, and as expected, the relative levels were consistent with the lower content in surface proteins in each kind of EVs.

**Figure 7.**
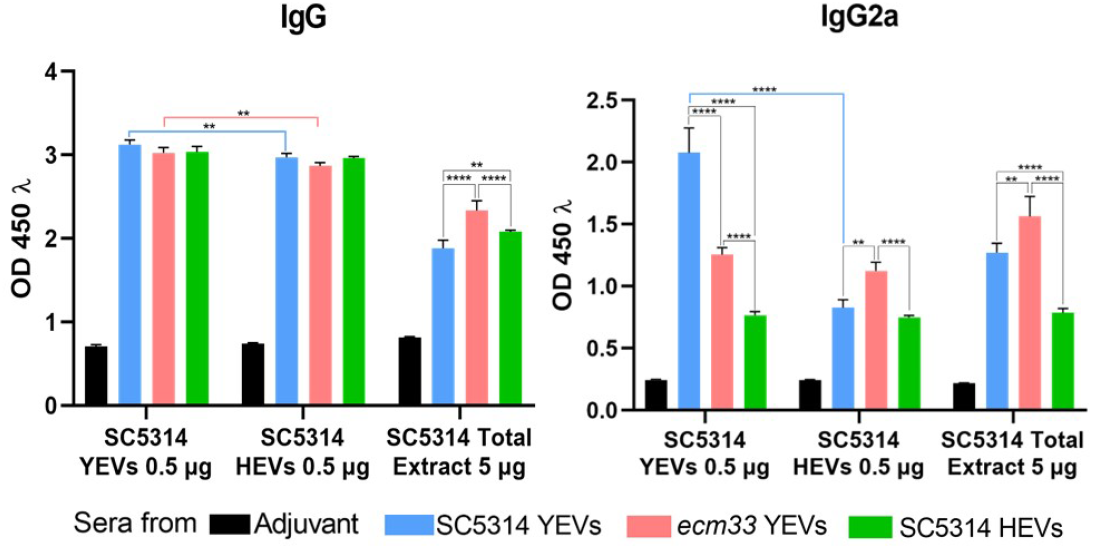
IgG and IgG2a antibody titers. ELISAs with pooled sera obtained prior to infection from immunized mice with SC5314 YEVs (blue), *ecm33* YEVs (pink), and SC5314 HEVs (green). Protein samples from SC5314 YEVs, SC5314 HEVs, and total cytoplasmic extracts from the wild-type strain SC5314 were used for detection purposes. Statistically significant differences in antibody titers were observed across all groups compared to the control group. Two-way ANOVA test * p <0.05, *** p <0.001, **** p <0.0001

Concerning the IgG2a antibody titers, each sera rendered different proportions of IgG2a against each extract. The best results were obtained with the sera from mice immunized with SC5314 YEVs against SC5314 YEVs protein sample, that rendered approx. 75% of IgG2a of the total IgG levels (Figure 7), in line with the highest protection against IC exhibited by these SC5314 YEVs. The other sera presented levels of IgG2a against the same YEVs extracts according to their relative level of protection.

The IgG2a titers in SC5314 YEVs sera dropped to less than a half against the other protein samples, indicating that protective proteins are less represented in them (Figure 7). The serum of mice vaccinated with *ecm33* YEVs contained the highest levels of IgG2a against the total cytoplasmic extract and similar levels against the HEVs and SC5314 YEVs proteins. These differences in IgG2a levels are likely attributed to the abundance in cytoplasmic proteins contained within these vesicles. In correlation with the diminished protective efficacy demonstrated by the S5314 HEVs, the sera from mice immunized with these vesicles consistently displayed the lowest levels of IgG2a across all protein samples. Abundance in proteins already described as immunogenic in each kind of EVs will be discussed later.

### 2.5. Analysis of cytokines present in the sera of immunized and surviving mice

We determined the cytokine profile for each of the pooled sera obtained at two time points indicated in figure 4: after complete immunization with the different EVs at the day prior to infection, and 30 days after infection. The measured cytokines were TNF-α, IFN-γ, IL-4, IL-10, and IL-17 (Figure 8). The lower levels were detected for cytokines involved in Th2 response: IL-10 was not detected in any sera and IL-4 was not detected in the surviving mice. For the other cytokines we found significant differences among the different groups of sera obtained before infection, as well as between the sera groups from immunized mice and the corresponding 30-day survivors for almost each cytokine.

**Figure 8.**
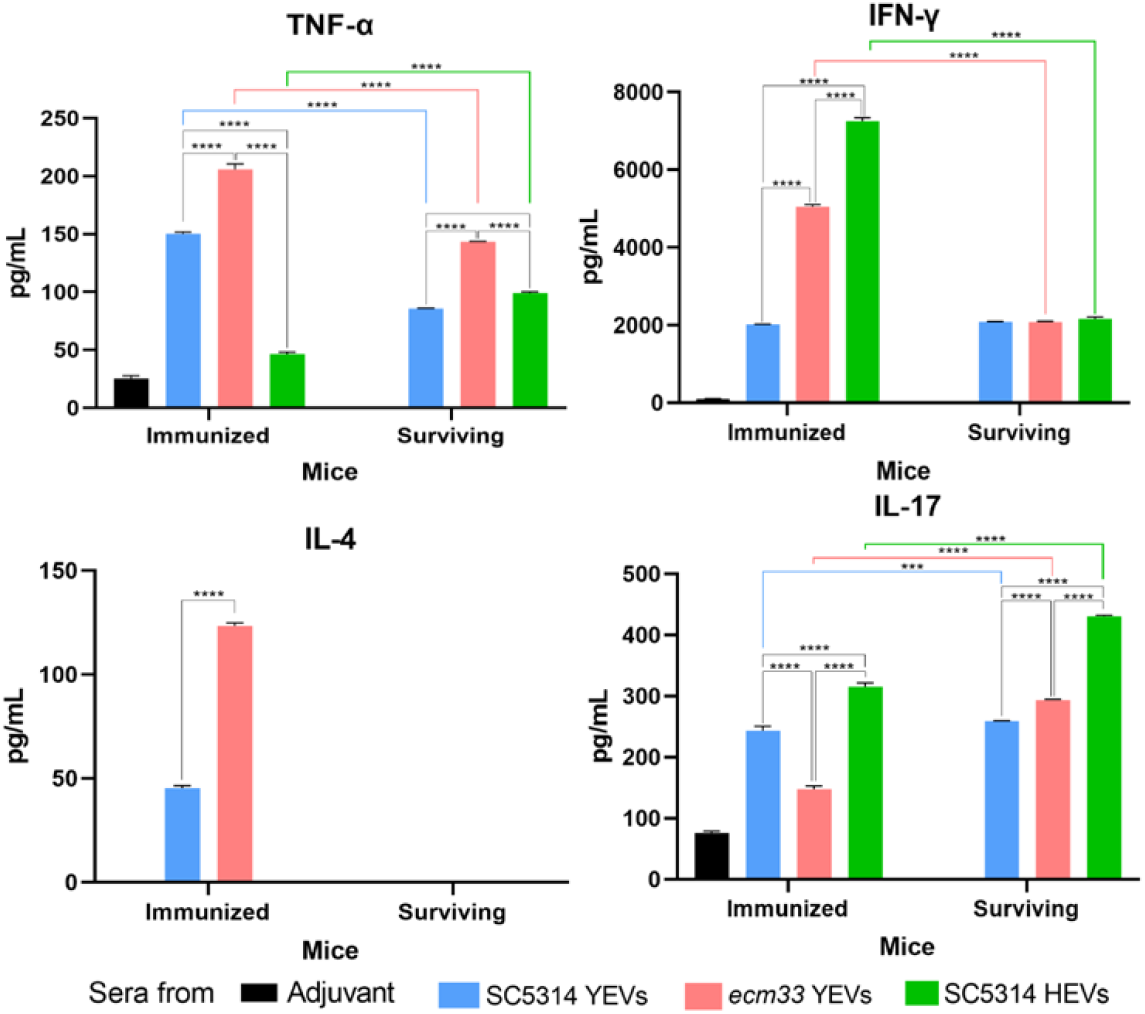
Cytokine profile for each of the pooled sera obtained after complete immunization with the different EVs (SC5314 YEVs (blue), *ecm33* YEVs (pink), and SC5314 HEVs (green)) but prior to infection (immunized), and 30 days after infection (surviving). Adjuvant group (control group) received only the adjuvant. Statistically significant differences in cytokine levels were observed across all cases compared to the control group. Two-way ANOVA test * p <0.05, *** p <0.001, **** p <0.0001

Concerning TNF-α and IFN-γ levels, associated with the inflammatory response, both were detected in sera from immunized and surviving mice. For TNF-α, the lowest levels (50 pg/mL) were obtained in the HEVs group before infection. This value doubled in the SC5314 YEVs group and was 4-times higher in the *ecm33* YEVs group. In contrast, in the surviving mice, the levels of this cytokine significantly decreased (½) in both the SC5314 YEVs and *ecm33* YEVs groups, while the opposite (2x) took place for HEVs surviving mice. *ecm33*Interestingly, IFN-γ values had an inverse relationship with the level of protection achieved by immunization with the different EVs, while the IFN-γ levels became comparable (2000 pg/mL) for the three groups of surviving mice.

Concerning IL-17, among immunized mice, the *ecm33* YEVs group had the lowest level (∼ 150 pg/mL), followed by the SC5314 YEVs group (250 pg/mL), and the HEVs group had the highest level (315 pg/mL). For surviving mice, compared to the immunized mice, a significant increase was observed for the *ecm33* YEVs and HEVs groups, while for the surviving SC5314 YEVs group, the mice with the highest level of protection, the increase in IL-17, although statistically significant, was very small.

## 3. Discussion

Several studies on *C. albicans* fungal EVs demonstrated that the protein content of EVs varies significantly depending on the strain and culture conditions [20, 35, 42, 43]. This variation in protein content is a crucial factor that can determine the role of EVs in providing protection or pathogenic effects (nicely reviewed in [44]). Here we compared the proteomic composition of EVs secreted from cells of the *C. albicans ecm33Δ* mutant and its parental strain and tested the protection ability against IC of these two types of EVs, as well as of SC5314 HEV.

Regarding *ecm33* YEVs, it’
ss important to note that the administration of the mutant as a live-cell vaccine previously demonstrated high protection against the virulent parental strain (87,5%) [11], whereas the immunization with 5 µg of *ecm33* YEVs with aluminum adjuvant resulted in protecting 64% of the mice infected with 5 ***x*** 10^5^ cells/mL of the parental strain SC5314. This level of protection is lower than that obtained with the mutant but it is in agreement with the fact that cellular vaccines with avirulent mutants usually produce a greater protective effect [9, 12, 45]. However, it represents a high level of protection compared to other acellular vaccines which are generally composed of proteins tested against IC. For example, the intravenous administration of the recombinant protein Als1 or Als3, together with Freund’s complete adjuvant, achieved around 40% protection against an intravenously (i.v.) infection with the same infectious dose like that used in our study [46].

We obtained an even higher level of protection against IC when using the SC5314 YEVs following the same immunization protocol, with 90,9% survival at the end of the experiment. Since the *ecm33Δ* mutant has an altered cell wall structure and different proteins exposed in the surface [11], it was logical to expect a different protein composition in the vesicles secreted by this mutant. Accordingly, in the label-free proteomic analysis of the *ecm33* YEVs vs the SC5314 YEVs, we observed a higher proportion of cell surface proteins in SC5314 YEVs. As these surface proteins are the initially exposed to the host immune system, antibodies against them are of great interest for achieving protection. For example, we identified Sap2, a major secreted aspartyl proteinase, exclusively in the SC5314 YEVs (Table 1). In a previous study, vaccination with this protein conferred immunoprotection in a rat model of vaginal candidiasis, eliciting specific anti-Sap antibodies of the IgG and IgA classes [47]. Furthermore, in our comparative analysis of the top 100 most abundant proteins in each type of vesicle used for immunization, only SC5314 YEVs contained the immunogenic proteins Mp65, Mdh1, and Ecm33. Mp65 is a cell surface mannoprotein highly immunogenic that induces extensive T-cell proliferation of human peripheral blood mononuclear cells and is a primary target of the human T-cell response to *C. albican* [48]. The protective capacity of the Mdh1p against IC has already been demonstrated by Shibasaki and colleagues after administration of this protein both by subcutaneous and intranasal route [49]. Ecm33 also plays a crucial role in *C. albicans* pathogenesis since the *ecm33Δ* mutant is completely avirulent [50]. Moreover, preliminary tests conducted by our research group, employing the identical immunization protocol utilized in this study, demonstrated that 5 µg of a recombinant Ecm33 protein can postpone the mortality of infected mice. (5***x***10^5^ cells/mL of SC5314 via i.v.) [51]. It is reasonable to conclude that the enhanced protective efficacy of SC5314 YEVs can be attributed, among other factors such as potential differences in lipid composition, to the presence of these immunogenic surface proteins.

To our knowledge, the 90.9% of protection achieved following vaccination with SC5314 YEVs against an acute and rapid *C. albicans* iv-induced candidemia with an infection dose of 5 ***x***10^5^ cells/mL, represents one of the most effective reports on the use of acellular vaccines. Previous studies have demonstrated a higher level of protection (100%) [36]; however, these studies used a different infection model—an intraperitoneal model where the control mice survived for 15 days. In vaccination assays with Mdh1, a high rate of survival was also reported. However, in that study, despite using an i.v. infection route, a lower dose of 1,1 ***x***10^5^ cells SC5314 cells was administered, which required nearly 25 days to cause death in the control mice [49]. In contrast, our model uses a higher dose for i.v.-induced candidemia, resulting in 100% mortality in a much shorter time frame. This intravenous challenge model has been widely employed to study the virulence of different strains, as it enables the fungus to reach internal organs, such as the kidneys, liver, and brain, mimicking parenteral infections that can occur via catheters in hospitalized patients. This route necessitates a rapid immune response to effectively control the infection. Although the high protection observed with the SC5314 YEVs is a very relevant result, we also obtained unexpected and important findings with the SC5314 HEVs, which appear to exhibit dual behavior. On the one hand, the SC5314 HEVs induced detrimental effects, resulting in the death of 63% of the mice days before those in the control group. On the other hand, 36.4% of the mice manage to overcome this phase, and a protective effect was subsequently observed. To understand this paradoxical outcome, it is important to note that some studies have linked fungal EVs to increased pathogenicity [52] or pro-inflammatory effects [53], while others have demonstrate immunoprotective effects [54]. The harmful effect of the SC5314 HEVs is consistent with the cytotoxicity observed *in vitro* on THP1 human macrophages exposed to these EVs [35]. Moreover, we can at least partly explain the dual behavior of the SC5314 HEVs based on the results of the comparative analysis of the most abundant proteins in the three types of EVs. Only SC5314 HEVs contained 7 proteins linked to pathogenesis among the top 100 most abundant proteins. Of particular interest, the candidalisyn precursor protein Ece1 was among them, which may contribute to the detrimental effect mentioned. On the other hand, the protective effect observed in 36% of the mice can also be explained by our proteomics results, which show that all three types of vesicles share several proteins already described as immunogenic and protective among their top 100 most abundant proteins. These include the moonlighting proteins Fba1, Pgk1, Cdc19 [23] and Bgl2 [20]. These findings align with previous literature describing the complex role of fungal EVs in pathogen-host interactions. Fungal EVs can act as mediators in cellular communication, delivering virulence factors and promoting infection, damage and inflammatory responses. However, EVs can also have protective effects, as they can inhibit fungal growth and stimulate adaptive immune responses.

Cytokines also play a crucial role in the resolution of candidiasis, orchestrating both innate and adaptive immune responses necessary to control and eliminate *Candida* cells. In our analysis of cytokines from the vaccination experiments, we were unable to detect IL-10, and IL-4 levels were very low, being only present in the sera of immunized mice and not in the 30-day survivors. These results suggest a weakened Th2 response. However, although cellular immunity plays a more critical role than humoral immunity in the defense against fungal infections [10], antibodies contribute to enhancing phagocyte activity and protection against IC. The use of aluminum-based adjuvant favors the induction of humoral responses [55]. In our experiments, total IgG titers were similar across all sera when tested against any of the EVs protein extracts, but lower when using total cell extract. This is consistent with the underrepresentation of cell wall and surface immunogenic proteins in cytoplasmic extracts. In fact, different strategies, such as cell fractionation or cell shaving, have to be used to obtain a high proportion of *C. albicans* cell surface proteins [56-59]. However, we observed clear differences in IgG2a titers depending on the serum and the protein extract used. The IgG2a subclass plays a key role in opsonization and activation of the complement system [60, 61] and has been associated with a protective effect against candidiasis [10, 62]. The highest IgG2a titers were observed in the serum from SC5314 YEVs-vaccinated mice against the protein extract from the same EVs. This group also exhibited the highest survival rate, confirming that a robust IgG2a response correlates with increased survival and better protection against IC.

The Th1 response and IFN-γ production are crucial for the activation of neutrophils and macrophages to eliminate *C. albicans* [63]. In fact, mice lacking the IFN-γ receptor are more susceptible to systemic candidiasis than IL-4 knock out (KO) mice [9]. Therefore, an effective vaccine should promote this Th1 response. The most advanced vaccine currently developed against systemic candidiasis is based on Als3p and generates specific T cells that produce IFN-γ and IL-17A, both of which are required for a protective response [64]. Our data on the cytokine levels align with the importance of the Th1 response, as all three vaccine formulations induced low levels of IL-4 and undetectable levels of IL-10, while high levels of IFN-γ in immunized mice and reminded detectable in the surviving mice even 30 days after infection.

Although a substantial volume of literature focuses on IL-17 role in oropharyngeal and mucocutaneous candidiasis [65-68], this cytokine is also implicated in systemic infections. IL-17 receptor (IL-17R) KO mice exhibited significantly reduced survival rates and higher kidney colonization during i.v. *C. albicans* infection, showing both the mobilization of peripheral neutrophils and their influx into infected organs significantly impaired and delayed [69]. In our study, immunization with SC5314 YEVs produced moderate IL-17 levels that showed minimal variation in surviving mice, while lower IL-17 levels were induced after *ecm33* YEVs inmmunization, correlating with the protection provided by each type of EV. The highest levels of this cytokine were found in the HEV-vaccinated sera of surviving mice, which increased compared to the pre-infection sera. Candidalysin has been associated with *Candida* virulence due to its cytolytic role [70]; however, a recent study also linked it to an immunoprotective function in fungal infections of the central nervous system [71]. In a recent review, this dual role of candidalysin was explained in relation to its quantity [72]. Under commensal conditions, minimal amounts of candidalysin are secreted, which may help maintain a balance between limited host damage and an effective immune response. In contrast, during infection, higher levels of candidalysin can damage host cells and tissues. The authors also propose an immunopathological state in which excessive candidalysin may trigger an overreaction of the immune system. As HEVs are the only vesicles containing the Ece1 precursor of candidalysin among their most abundant proteins, it would be interesting to further investigate a potential relation between the presence of this protein, the elevated levels of IL-17 and IFN-γ observed in mice vaccinated with this type of EVs and a potential exacerbated inflammatory response that may be detrimental to most of the mice.

In summary, our results demonstrate the potential of YEVs as vaccine candidates against IC. Additionally, a dual role in the host response, depending on the type of EVs used for the immunization protocol, was observed: YEVs are associated with protection, while HEVs either increase virulence or confer a protective effect in the mice. These align with the distinct roles of *C. albicans* cell morphologies in host-fungal interaction, with the yeast form associated with the commensal state and hyphae involved in pathogenesis.

## 4. Materials and Methods

### 4.1. Fungal strains

The *C. albicans* clinical isolate SC5314 (Gillum et al., 1984) and the cell wall *ecm33Δ* mutant strain RML2U (*ecm33Δ::hisG/ecm33Δ::hisG ura3Δ::imm434/ura3Δ::imm434::URA3*.) [39, 50] were used in this work.

### 4.2. Culture conditions and extracellular vesicles isolation

Extracellular vesicles (EVs) were isolated from yeast and hyphal forms of *C. albicans* as follows: EVs were derived from the yeast forms of two *C. albicans* strains: the parental strain SC5314, referred to as SC5314 YEVs and the *ecm33*Δ mutant strain, referred to as *ecm33* YEVs. EVs were also collected from the hyphal forms of the clinical isolate SC5314, referred to as SC5314 HEVs.

To isolate SC5314 YEVs or *ecm33* YEVs, a total of 1 L of YPD culture media for each strain were utilized, incubated for 16 h at 37°C and 180 rpm. To isolate SC5314 HEVs the hyphal cells were grown in YNBS culture media supplemented with 75 mM of 3-(N-morpholino) propanesulfonic acid (MOPS and 5 mM of N-acetylglucosamine (N-AcGlc) adjusted to a pH of 7.4.

The procedure for isolating EVs was carried out following the methodology described by Gil Bona et al. [37]. All the steps of the process were performed at 4°C. Briefly, supernatants obtained from the corresponding culture media were collected by centrifugation at 8,000 rpm for 40 min in a Beckman Coulter J2-HS centrifuge equipped with the JA-10 rotor. The collected supernatants were then filtered through a 0.45 µm filter to remove any remaining cells and debris to ensure the purity of the EVs samples. To preserve the EVs integrity, one protease-inhibitor tablet (Pierce™ EDTA-free, Thermo Fisher) and 1 mL of Phenylmethanesulfonyl fluoride (PMSF) were added to each litre of filtered supernatant. The supernatants containing the EVs were concentrated using a Centricon Plus-70 filter with a cutoff of 100 kDa (Millipore). This concentration step was achieved by centrifuging the samples at 3,500 rpm in an Eppendorf 5810R centrifuge, resulting in a final volume of 8 mL. The concentrated supernatants were then subjected to ultra-centrifugation at 100,000 g for 1 h in a Beckman Optima XL-90 centrifuge with a 90 Ti rotor. As a result, the EVs formed pellets at the bottom of the tubes. The pellets containing the isolated EVs were washed twice with PBS (phosphate-buffered saline) to remove any impurities or contaminants and then solubilized in 50 µL of sterile PBS. The protein concentration of the isolated EVs samples was determined using the Bradford assay with a reactive solution from BioRad, following the instructions provided by the manufacturer. Fresh preparations of the EVs suspension were kept at 4 °C until used for the immunization experiments.

### 4.3. Transmission electron microscopy

Samples for transmission electron microscopy were prepared according to the protocol described by Miret et al. [73]. Briefly, EVs were fixed at 4°C for 24 h in 500 µl fixative solution (sodium cacodylate buffer, pH 7.2, containing 1.2% glutaraldehyde and 2% paraformaldehyde). The samples were then washed with saline and postfixed for 90 min with 1% potassium permanganate. The fixed EVs were dehydrated through a graded series of ethanol and embedded in Embed 812 resin (Electron Microscopy Sciences, Hatfield, PA). Thin sections were stained with uranyl acetate and lead citrate and then imaged with a Zeiss EM902 electron microscope. Multiple fields containing numerous EVs were imaged.

### 4.4 NTA Analysis

Nanoparticle Tracking Analysis (NTA) was performed using a Zeta-View instrument (Particle Metrix). The samples were diluted in prefiltered PBS to approximately 106 to 107 particles/mL. A CMOS camera with 405 nm 68 mW laser was used to scan 11 cell positions capturing 30 frames per position at 25°C with camera sensitivity 85, shutter speed 100. Data analysis was conducted with ZetaView Software (version 8.05.12 SP1), applying settings for minimum brightness of 30, maximum brightness of 255, minimum area of 5, maximum area of 1,000, and minimum trace length of 15. Each sample was recorded in triplicate in light scatter mode. Particle size and concentration were subsequently analyzed and visualized using GraphPad Prism 8.0 software.

### 4.5. Proteomic studies

#### 4.5.1. Digestion and desalting of peptides

50 µg of proteins were separated by SDS-PAGE in a mini-protean system (BioRad), (with a 4 cm resolving gel of 10% acrylamide (acrylamide, 19:1) in 1.5 M Tris, pH 8.8, and a 4 cm stacking gel of 4% acrylamide in 0.5 M Tris, pH 6.5 V) until the electrophoresis front (bromophenol blue) advanced about 2 cm into the stacking gel. The protein bands from the stacking gel were excised, and disulfide bonds were reduced with 10 mM DTT in 25 mM AMBI at 56°C for 30 min and then alkylated with 22.5 mM iodoacetamide in 25 mM AMBI for 10 minutes in the dark. Digestion was carried out with recombinant proteomics-grade trypsin (Roche) at a ratio of 1:50 in 50 mM AMBI and incubated overnight at 37°C. Peptides were collected from the digestion supernatant and extracted from the gel with ACN. The resulting solution was dried in a Speed-Vac and reconstituted in 12 µL of 2% ACN, 0.1% formic acid. Peptide concentration was measured by fluorimetry using the Qubit 3.0 system from Thermo Fisher Scientific. From the digested peptide mixture, 0.5 µg of the sample were injected into the Vanquish Neo IFC chromatograph (Thermo), concentrated in a PEPMAP100 C18 NanoViper Trap pre-column (Thermo Fisher).

#### 4.5.2. LC-MS/MS

The peptides were separated by chromatography in a 50 cm PEPMAP RSLC C18 column (Thermo) with a gradient from 2% to 35% ACN, 0.1% formic acid over 90 min before analysis in the mass spectrometer. The peptides were then ionized by electrospray in positive mode and analyzed in a Q Exactive HF mass spectrometer (Thermo) using DDA (data-dependent acquisition) mode. From each MS scan (between 350 and 1800 Da), the 15 most intense precursors (charge between 2+ and 5+) were selected for HCD (high collision energy dissociation) fragmentation, and the corresponding MS/MS spectra were acquired.

#### 4.5.3. Shotgun Protein Identification

The data files generated from the shotgun analysis were transferred to the Proteome Discoverer software (Thermo), using MASCOT v2.6 as search engine on the Candida Genome Database (6209 sequences). The following parameters were used for the searches: tryptic cleavage, up to two missed cleavage sites allowed, and tolerances of 10 ppm for precursor ions and 0.02 Da for MS/MS fragment ions and the searches were performed allowing optional Methionine oxidation and fixed carbamidomethylation of Cysteine. A search was also performed against a database containing common contaminant proteins. When a peptide could be assigned to multiple proteins, the software applied the principle of parsimony to generate a “Master” protein, which was reported in the results. The Percolator algorithm was used to validate identified proteins with a 1% FDR. The mass spectrometry proteomics data have been deposited to the ProteomeXchange Consortium via the PRIDE (Perez-Riverol et al., 2022) partner repository with the data set PXD055891 and 10.6019/PXD055891.

#### 4.5.4. Protein quantification

To determine the abundance of the identified proteins in the samples, a label-free experiment was performed. The processing workflow was initiated with an alignment of the chromatograms of all the samples with a tolerance of up to 10 min. Subsequently, the quantification of the precursor was based on ions signal intensity of the unique peptides. Finally, the results were normalised to the total amount of the peptides, equalling the total abundance among the different samples. The protein quantification design using Proteome Discoverer software v2.4 (Thermo) was not paired between each group with a pairwise ratio approach calculation. The Proteome Discoverer includes a statistical feature (Anova Background) for assessing the significance of differential expression by providing P values for those ratios. Only proteins identified with high confidence (FDR < 1%) with at least two unique peptides were considered as differentially expressed between groups.

### 4.6. Mouse model

Groups of 11 inbred female BALB/c mice, aged 6-8 weeks and weighing approximately 20g, (obtained from Harlan France) were used for the immunization experiments, while the control groups had only 8 mice. The mice were immunized twice, with a two-week interval between immunizations, by subcutaneously injecting 100 µL of the respective EVs suspension (the specified amount of protein for each case is detailed in result section and the Figures S1, 3 and 4) or PBS, plus adjuvant if required. The EVs suspensions used were SC5314 YEVs, *ecm33* EVs, or SC4514 HEVs, in PBS. In the experiments with adjuvant, the EVs suspension was formulated with Imject TMAluminum adjuvant (Thermo), in a 1:2 proportion for the first experiment and 1:3 (v/v) for the subsequent experiments. For the evaluation of the protection elicited by alive *ecm33* mutant cells, a dose of 2.5 × 106 cells of the *ecm33Δ* mutant was prepared in a sterile saline solution and administered i.v. via the caudal vein as previously described (Martinez-Lopez et al., 2008). Fifteen days after the second immunization, all mice were infected intravenously through the tail vein with a lethal dose of 1 × 106 or 5 × 10^5^ cells from the wild-type strain SC5314. *C. albicans* inoculum for infection was grown in a shaking incubator for 14 h at 30°C in YPD medium (2% glucose, 0.5% yeast extract, 0.5% peptone extract). Yeast cells were harvested, washed twice with sterile, apyrogenic PBS, counted in a hemocytometer, and suspended at the appropriate concentration. Blood samples were collected from 2 mice in each group on day 15 after the second immunization and prior to infection. Additionally, blood samples were collected from all surviving mice 30 d after the lethal infection. The blood samples were incubated at 37º for 1 h and subsequently centrifuged at 10,000 x g for 10 min, and the serum (upper phase) was collected and stored at −80 °C for further analysis. Throughout the experiment, all mice were treated in accordance with the ethical guidelines for animal experimentation, as specified in Protocol PROEX 088/77. The number of deaths was recorded daily, and survival curves were plotted. Statistical analysis was performed using the log-rank (Mantel-Cox) survival test.

### 4.7. Preparation of protein cytoplasmic extracts

Cells of the SC5314 strain were harvested and subsequently washed twice in cold PBS. The cells were then resuspended in a lysis buffer consisting of 50 mM Tris-HCl (pH 7.5), 1 mM EDTA, 150 mM NaCl, 1 mM DTT, 0.5 mM PMSF, and 10% of a mixture of protease inhibitors (Pierce). Subsequently, the cells were disrupted by the addition of glass beads (0.4-to 0.6-mm diameter) using a Fast-Prep system (Bio101; Savant), applying 5 cycles of 20 seconds each with 1 minute of cooling on ice between cycles. The resulting cell extracts were separated from the glass beads through centrifugation, and the supernatant was collected. To ensure clarity, the supernatant was further cleared by centrifugation at 13,000 rpm for 15 min at 4°C. Finally, the protein concentration of the obtained supernatant was determined using the Bradford assay

### 4.8. Measurement of antibody titers

The total titer of IgG and IgG2a was determined using three different protein samples: total cytoplasmic protein extract (TPE SC5314), and proteins from SC5314 YEVs and SC5314 HEVs, both treated with 1% ASB-14 detergent (Sigma-Aldrich) in PBS buffer with 0.1% BSA. The 96-well plates were coated with these protein samples in triplicates and incubated overnight at 4°C. Serum pools from immunized mice with each of the different types of EVs were tested in triplicate. The next day, the plates were washed twice with PBS containing 0.05% tween-20 and 0.1% BSA to remove any unbound material and were blocked with a 5% power milk solution for 2 h at 37°C to prevent non-specific binding. After another round of washing, sera from the different groups were added to the plates in triplicates for each type of protein sample in 1:1000 dilution in PBS with 1% BSA and incubated overnight at 4°C. Following incubation, the plates were washed again, and 50 µLof Invitrogen peroxidase-labeled anti-mouse IgG (Cat # 32430) or IgG2a (Cat # M32207), diluted 1:5000 in PBS with 1% BSA, were added and incubated for 2 h at room temperature. After washing the plates 5 times with the washing buffer, a TMB (Tetramethylbenzidine) substrate BD OptEIA™ was added. Color development was allowed for 5-10 min, and the reaction was stopped by adding 1 N sulfuric acid. The optical density was measured at 450 nm (OD450). The measured OD450 values were converted to antibody titers by calculating the reciprocal of the highest dilution factor that resulted in an OD450 above the cut-off threshold, which was determined as the average OD450 of blank wells plus 2 standard deviations.

### 4.9. Cytokines Measurement

The concentrations of cytokines TNF-α (Cat#BM607-3), IL-4 (Cat#BM613), IL-10 (Cat#BMS614), IL-17 (Cat#BMS6001), and IFN-γ (Cat#BM609, ExtraSensitive) were measured in the different sera according to manufacturer’s instructions by ELISA kits (Invitrogen). Briefly, TNF-α, IL-4, IL-10, IL-17, and IFN-γ were measured using 96-well plates coated with specific capture antibodies for each cytokine. The plates were incubated at 4°C overnight, or at room temperature for 2 hours, according to the kit protocol. After removing the coating solution, non-specific binding was blocked by adding 200 µL of blocking buffer (2% BSA in wash buffer) to each well, followed by incubation for 1 hour at room temperature. Serum samples, diluted 1:100 in PBS with 1% BSA, and known standards were added to the wells (100 µL per well) and incubated for 1 hour at room temperature. After washing, 100 µL of biotinylated detection antibody, specific to the target cytokine, was added to each well and incubated for 1 hour. Subsequently, 100 µL of streptavidin-HRP conjugate was added to each well, followed by another 1-hour incubation. The colorimetric reaction was initiated by adding 100 µL of TMB, which reacted with the HRP enzyme to produce a color change. The intensity of the color change was measured using a microplate reader at a wavelength of 450 nm, and cytokine concentrations were determined by comparing the optical density of the samples with a standard curve generated from known concentrations of the cytokines. Measurements were done in triplicates.

### 4.10. Quantification and Statistical Analysis

Log-rank test (Mantel-Cox) was used for survival data analysis. For parametric data, 2 groups were analyzed by two-tailed Student’s t-test and multiple groups were analyzed by analysis of variance (ANOVA). Sidak correction was performed, and the statistical significance was set at a P-value of 0.05, 0.001 or 0.0001

## 5. Conclusions

The different EVs derived from C. albicans elicit distinct effects on the host’s immune system, with varying levels of protection against subsequent systemic *C. albicans* infection. Notably, yeast-derived EVs, especially wild-type YEVs, offer significant protection compared to HEVs. Although the *ecm33Δ* mutant provided high protection when used as live cells, the EVs secreted by this mutant offered less protection than YEVs from the wild-type strain, but better than HEVs. This differential immune response appears to be closely linked to the protein composition of these EVs, with those containing a higher proportion of cell surface and specific highly immunogenic proteins offering stronger protection. In addition, our results revealed harmful effects from immunization with HEVs. All these results highlight the importance of carefully optimizing the composition of EVs. Analysis of immunized mice sera has shown that IgG2a antibody titers correlate with protection levels. Furthermore, variations in Th1 and Th17 responses depend on the type of EVs used in immunization. The complexity of these immune responses highlights the need for further research to develop an effective vaccine for IC.

## Supplementary Materials

The following supporting information can be downloaded at: www.mdpi.com/xxx/

Figure S1: Vaccination schedule (a) and survival curves (b) of a mouse model of invasive candidiasis showing the protective effect of immunization with different *C. albicans* EVs. Mice were immunized with different doses of EVs with adjuvant (Ad) or without adjuvant and subsequently challenged with an intravenous lethal dose of SC5314 (1*x*10^6^ cells). Statistically differences between immunization and control groups are marked with an asterisk. * Mantel-Cox test p <0.05;

Table S1: Proteins identified in SC5314 YEVs; Table S2: Proteins identified in *ecm33* YEVs; Table S3: Quantified proteins in SC5314 YEVs and *ecm33* YEVs; Table S4: List of proteins that significantly changed their abundance between SC5314 YEVs and *ecm33* YEVs

## Author Contributions

Conceptualization, R.ML., G.M., C.PG., C.G. and L.M.; methodology, R.ML., G.M., C.PG., M.C., C.G. and L.M.; validation, R.ML, C.PG. and ML. H.; formal analysis, R.M, G.C and M.L.; investigation, R.ML., G.M., C.PG., M.C., C.GD., G.C., F.C, and ML. H.; resources, C.G. and L.M.; data curation, R. ML. and M.C.; writing—original draft preparation, R. ML; writing—review and editing, R. ML., G.M. and L.M.; visualization, R. ML. and G.C.; supervision C.G and L.M.; project administration, C.G..; funding acquisition, C.G and L.M.. All authors have read and agreed to the published version of the manuscript.

## Funding

This research was funded by Spanish Ministry of Science and Innovation/ State Research Agency 10.13039/501100011033, grant number PID2021-124062NB-I00

## Institutional Review Board Statement

The study was conducted in accordance with the Declaration of Helsinki and approved by the Institutional Review Board (or Ethics Committee) of Consejería de Medio Ambiente, Administración local y Ordenación del Territorio de la Comunidad de Madrid (protocol code PROEX 088/77 on 20 of November of 2017).

## Data Availability Statement

The mass spectrometry proteomics data have been deposited to the ProteomeXchange Consortium via the PRIDE (Perez-Riverol et al., 2022) partner repository with the data set PXD055891 and 10.6019/PXD055891.

## Acknowledgments

The proteomic analyses were carried out at the Biological Techniques CAI, specifically within the Proteomics Unit at the Complutense University of Madrid (UCM). Similarly, the electron microscopy analyses were conducted at the UCM ICTS CNME ELECMI - National Center for Electron Microscopy. We acknowledge Dr. R. Prados-Rosales at laboratory at the School of Medicine, Universidad Autónoma de Madrid, Spain, for his help with NTA analyses. Guillermo Castejón enjoyed a contract under the Investigo program, within the framework of the Recovery, Transformation, and Resilience Plan, by the Ministry of Labor and Social Economy and the State Public Employment Service, regulated by Ministerial Order TES/1267/2021, assigned to reference CT19/23-INVM-65. Claudia Parra Giraldo enjoyed a Maria Zambrano contract from the Ministry of Science, Innovation, and Universities.

## Conflicts of Interest

The authors declare no conflicts of interest

## Notes

https://www.ebi.ac.uk/pride/

### Competing Interest Statement

The authors have declared no competing interest.

