## Supplementary figures and images for "From high protection to lethal effect: diverse outcomes of immunization against invasive candidiasis with different *Candida albicans* extracellular vesicles"

### Supplemental Figure 1

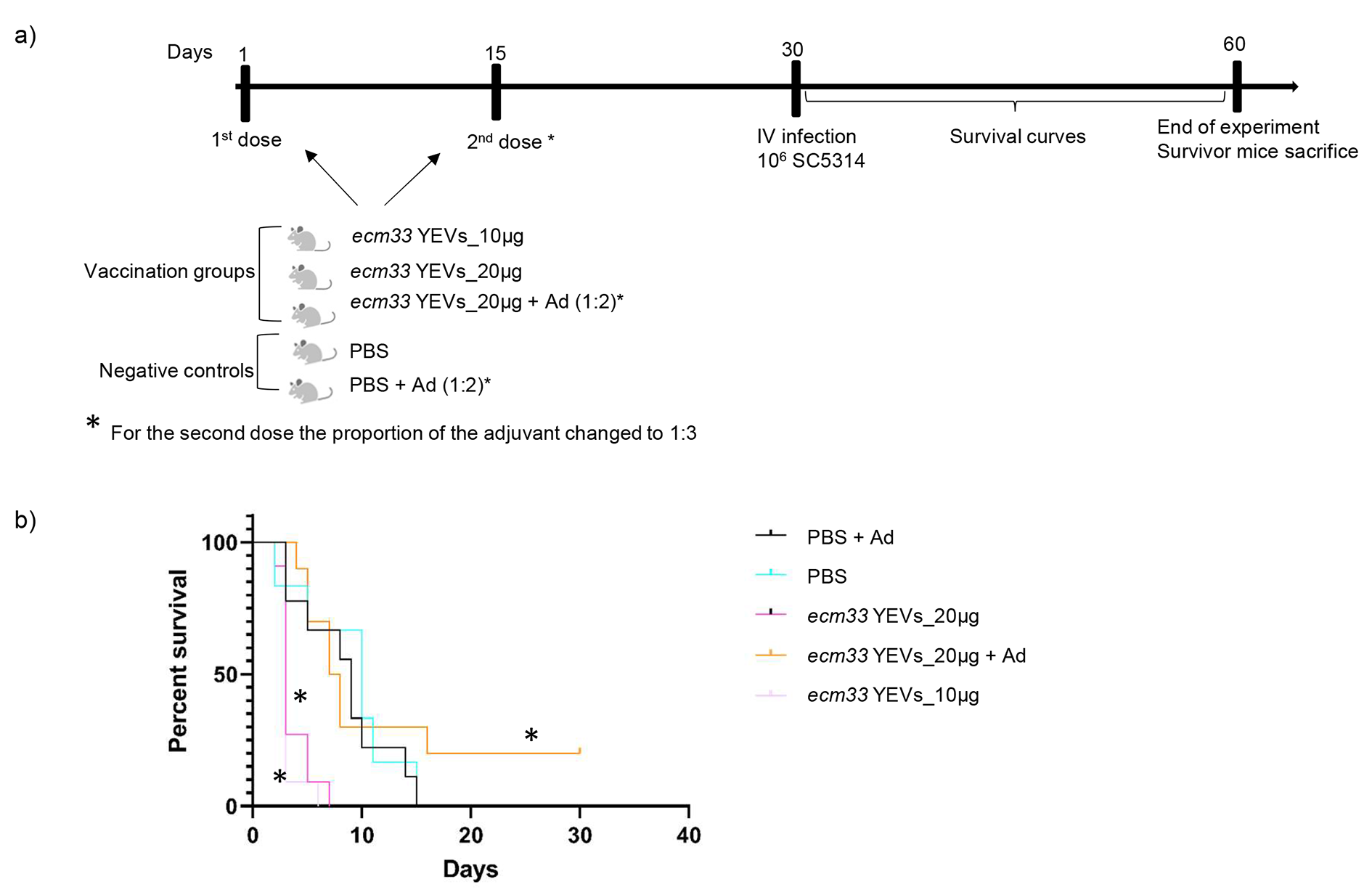
